# Differential effects of prolonged post-fixation on immunohistochemical and histochemical staining for postmortem human brains

**DOI:** 10.1101/2024.05.13.593904

**Authors:** Weiya Ma, Eve-Marie Frigon, Josefina Maranzano, Yashar Zeighami, Mahsa Dadar

## Abstract

Immunohistochemical (IHC) and histochemical (HC) staining is widely used for human brains post-fixed in formalin and stored in brain banks worldwide for months, years and decades. Understanding the effect of prolonged post-fixation, postmortem interval (PMI) and age on these staining procedures is important for interpreting their outcomes, thus improving diagnosis and research of brain disorders afflicting millions of world populations. In this study, we performed both IHC and HC staining for prefrontal cortex (PFC) of postmortem human brains post-fixed for 1, 5, 10, 15 and 20 years. A negative correlation was detected between the intensity of neuron marker neuron nuclear specific marker (NeuN), microglia marker ionized calcium-binding adaptor molecule 1 (Iba1), cresyl violet (CV) and Luxol fast blue (LFB) staining versus post-fixation durations. By contrast, a positive correlation was seen between the intensity of astrocyte marker glial fibrillary acidic protein (GFAP) and hemaetoxylin and eosin Y (H&E) staining versus post-fixation durations. No correlation was found between the staining intensity of NeuN, GFAP, Iba1, H&E, CV and LFB versus PMI. Moreover, no correlation was seen between NeuN, Iba1, H&E, CV and LFB, except GFAP, versus age. These data suggest that prolonged post-fixation exerts both positive and negative effects, but age and PMI have limited effects, on these IHC and HC parameters. Hence these differential changes need to be considered in interpretation of the results when using tissues with prolonged post-fixation. Furthermore, if feasible, it is recommended to perform IHC and HC staining for human brains with the same post-fixation time windows and to use the most optimal antibodies to offset its impact on downstream analyses.

## 1. Introduction

Immunohistochemistry (IHC) and histochemistry (HC) are widely used for diagnostic and research purposes. Both approaches are essential to brain research exploring the mechanisms underlying normal brain functions as well as neurological and psychiatric disorders. Postmortem human brains are important research materials to uncover the mechanisms underlying brain-related disorders. In brain banks worldwide, it is a common practice to postfix the donated postmortem human brains in 10% neutral-buffered formalin, i.e., 4% formaldehyde solution buffered to a neutral pH until future use. As such, most postmortem brains have usually been preserved in neutral formalin for months, years and decades (Vonsattel et al., 2008; Beach et al., 2015). An important concern in using these tissues is whether prolonged formalin post-fixation of postmortem brains affects the quality of subsequent IHC and HC staining. Although neutral formalin provides excellent tissue preservation to prevent degradation in IHC (Paavilainen et al., 2010), at the same time it leads to cross-linking of proteins and nucleic acids in tissues (Helander, 1994), thus masking antigen epitopes and reducing immunoreactivity in IHC (Arnold et al., 1996) and mRNA signals in *in situ* hybridization (Mostegl et al., 2011). Hence prolonged formalin post-fixation is generally believed to prevent the efficient use of postmortem human brains archived for years and decades. To improve IHC staining, some procedures including enzyme-based and heat-based antigen retrieval (HIAR) have been applied to break formalin-induced crosslinks in post-fixed tissues (Shi et al., 2011). Of these approaches, citrate buffer-based HIAR has widely been used for IHC staining on post-fixed human brains (Shi et al., 1998; Jiao et al., 1999; Ramos-Vara and Beissenherz, 2000; Leong et al., 2010). Using HIAR, the effect of prolonged post-fixation on IHC and/or HC staining for paraffin embedded human brain sections were explored (Lyck et al., 2008; Webster et al., 2009; Alrafiah and Alshali, 2019; Wu et al., 2022). These studies showed that the influence of prolonged post-fixation on IHC and/or HC staining was either negative or unaltered. However, the time spans of post-fixation for the examined group versus the control group from these prior studies were either very short (weeks or months) (Webster et al., 2009; Pikkarainen et al., 2010; Wu et al., 2022) or rather wide (a group with mixed post-fixation times from 1 to 20 years) (Alrafiah and Alshali, 2019). Furthermore, the detailed information on post-fixation times for presented data were missing and no quantitative analysis was performed (Alrafiah and Alshali, 2019), so it was not possible to systematically correlate the qualitative data with the post-fixation times.

In this study, we aimed to determine the effect of formalin post-fixation for 1, 5, 10, 15 and 20 years on IHC and HC staining for postmortem human brains. IHC staining for neuron nuclear specific marker (NeuN), astrocyte marker glial fibrillary acidic protein (GFAP) and microglia marker ionized calcium-binding adaptor molecule 1 (Iba1) are commonly used to evaluate the three major cell types in human brain tissues. NeuN, GFAP and Iba1 are important cell type specific markers involved in various brain functions and in pathogenesis of numerous brain disorders and widely used in brain research (Reddy and Abeygunaratne, 2022). Therefore, it is important to examine whether prolonged post-fixation exerts an effect on IHC staining of the three markers in human brains. Hematoxylin (H) and eosin Y (E) staining as well as cresyl violet (CV) staining are very common HC staining methods extensively used in clinical diagnostic and brain research. The two staining techniques help to identify different types of cells and tissues in the human brain and provide important information regarding the pattern, shape, and structure of cells. Hence it is also necessary to determine the effect of prolonged post-fixation on H&E- and CV-based HC staining for human brains.

Axonal myelination in both central and peripheral nervous system is essential for a fast and efficient transmission of electrical signals along neurons (Nave and Werner, 2014). Demyelination or myelin damage leads to numerous neurological conditions (Stadelmann et al., 2019), such as multiple sclerosis (Staugaitis et al., 2012). It also contributes to white matter lesions/hyperintensities in neurodegenerative diseases, including Alzheimer’s disease (Papuc and Rejdak, 2020). Myelin research on postmortem human brains is essential to understand the underlying mechanisms of these disorders. Luxol fast blue (LFB) staining is a most common HC technique to illustrate normal myelin as well as to assess myelin damage, and as an indirect marker of axonal degeneration in diagnosis and research of these diseases. Hence it is important to evaluate the effect of prolonged post-fixation duration on LFB myelin staining for human brains.

Postmortem interval (PMI) was defined as the time elapsed after death until the samples were fixed in formalin. It is generally believed that a shorter PMI could alleviate degradation of proteins and nucleic acids to preserve tissue and cell structures. Prior studies found that PMI over days did not seem to significantly affect IHC staining (Blair et al., 2016; Mizee et al., 2017; Krassner et al., 2023). Regarding the role of the donor age in IHC staining, mixed outcomes were reported (Gonzalez-Maeso et al., 2002; Mizee et al., 2017; Krassner et al., 2023). In this study, we also examined whether PMI and donor age play a role in IHC and HC staining for human brains.

To address the above-mentioned issues, in the current study we performed NeuN, GFAP and Iba1 IHC staining as well as H&E, CV and LFB HC staining in the free-floating or mounted cryosections of prefrontal cortex (PFC) from human brains post-fixed for 1, 5, 10, 15 and 20 years. We also performed a series of linear regression analysis to examine the relationships between IHC and HC staining versus post-fixation time, PMI and donor age.

## 2. Materials and methods

### 2.1 Postmortem human brain acquisition

All experiments were performed with the approval of Human Research Ethics Committees of McGill University and Douglas Mental Health University Institute. After reception of the whole brain at the Douglas-Bell Canada Brain Bank (DBCBB; https://douglasbrainbank.ca), postmortem human brains were collected and hemispheres were immediately separated by a sagittal cut in the middle of the brain, brainstem, and cerebellum. One unsliced hemisphere (right or left, in alternation), 1⁄2 brainstem, and 1⁄2 cerebellum were fixed in 10% neutral buffered formalin. For the current study, we requested PFC blocks (n=20) from human brains which have been post-fixed for 1, 5, 10, 15 and 20 years (n=4 per group, in total, n=20). All specimens were similarly processed and post-fixed in the same solution (i.e. 10% formalin), with the only difference in post-fixation times. All brains were donated to the DBCBB by familial consent through the Quebec Coroner’s Office. Blood toxicology assessment was performed and individuals with evidence of drugs or psychotropic medications were excluded. Individuals with a known history of neurological disorders or head injury were also excluded from this study. Demographic characteristics associated with each sample are listed in Table 1. Groups were matched as closely as possible for sex, age, race and PMI.

**Table 1.** Complete demographic for human brain cohort.

| Groups | Genders | Ages | Races | Causes of death | PMI (hours) | Post-fixation (years) |
| --- | --- | --- | --- | --- | --- | --- |
| 1yr-1 | M | 50 | Caucasian | Natural | 54.50 | 1 |
| 1yr-2 | F | 41 | Caucasian | Natural | 51.50 | 1 |
| 1yr-3 | M | 19 | Caucasian | Suicide | 46.00 | 1 |
| 1yr-4 | M | 33 | Caucasian | Accidental | 70.50 | 1 |
| 5yr-1 | M | 53 | Caucasian | Accidental | 49.50 | 5 |
| 5yr-2 | M | 59 | Caucasian | Suicide | 47.50 | 5 |
| 5yr-3 | M | 55 | Caucasian | Accidental | 51.00 | 5 |
| 5yr-4 | M | 59 | Caucasian | Natural | 46.42 | 5 |
| 10yr-1 | M | 75 | Caucasian | Natural | 15.88 | 10 |
| 10yr-2 | M | 91 | Caucasian | Natural | 72.68 | 10 |
| 10yr-3 | M | 80 | Caucasian | Natural | 42.58 | 10 |
| 10yr-4 | F | 84 | Caucasian | Natural | 10.72 | 10 |
| 15yr-1 | F | 49 | Caucasian | Suicide | 59.50 | 15 |
| 15yr-2 | M | 81 | Caucasian | Accidental | 98.75 | 15 |
| 15yr-3 | M | 30 | Caucasian | Suicide | 51.50 | 15 |
| 15yr-4 | M | 36 | African | Natural | 55.25 | 15 |
| 20yr-1 | F | 25 | Caucasian | Suicide | 20.00 | 20 |
| 20yr-2 | M | 22 | Caucasian | Suicide | 24.00 | 20 |
| 20yr-3 | F | 55 | Caucasian | Suicide | 36.00 | 20 |
| 20yr-4 | F | 44 | Caucasian | Suicide | 29.50 | 20 |

Footnotes: Age of the individuals (years): 19-91, average: 52.05; Postmortem interval (hours): 10.72-98.75, average: 46.66.

### 2.2 Cryostat sectioning of PFC samples

After reception from the DBCBB, 20 formalin-fixed PFC containing blocks from human brains post-fixed for 1, 5, 10, 15 and 20 years were transferred to 30% sucrose in 1X phosphate buffered saline (PBS) for cryoprotection until sectioned on cryostat. PFC blocks were embedded in M1 embedding medium (Fisher Scientific, Saint-Laurent, Quebec, Canada) and cut into 10- and 50-μm-thick sections (Leica, Feasterville, Pennsylvania, USA). The 10-µm-thick sections were cut from the middle part of the blocks and sequentially mounted on gelatin pre-coated slides and stored in −80°C freezer until they were used for HC such as H&E and CV staining. The 50-µm-thick sections were sequentially collected in cryoprotectant contained in 24-well culture plates and kept in −20°C until being used for IHC and HC such as LBF staining.

### 2.3 NeuN, GFAP and Iba1 immunohistochemistry

On the day of free-floating immunostaining, PFC sections were taken out of the freezer and washed in 1XPBS, then transferred into Eppendorf tubes containing the antigen retrieval buffer (10mM sodium citrate, 0.05% Tween 20, pH 6.0). The Eppendorf tubes were heated on boiling water (100 to 110°C) in a 1-liter beaker for 20 min. Then the tubes were cooled down on ice and sections were rinsed with cold double distilled (dd) H_2_O for 10 min. Sections were transferred into 1X PBS containing 0.05% Triton-100 (PBS+T) in wells of a 24-well culture plate. Sections were quenched in 0.03% H_2_O_2_ for 15 min to abolish the endogenous peroxidase in the tissue and next incubated in 50% ethanol for 1 hour to increase the tissue penetration by antisera. Sections were incubated in 10% normal goat serum (NGS) and 1% Bovine albumin diluted in PBS+T for 2 hours to block the endogenous non-specific antigenic sites. Finally, sections were incubated, respectively, in a mouse monoclonal antibody raised against a neuronal marker NeuN antibody (1:5000, Abcam, Cat. ab10424, Toronto, Ontario, Canada), in a rabbit monoclonal antibody raised against a neuron marker (1:5000, Abcam, ab177487), in a polyclonal rabbit antiserum raised against an astrocyte marker GFAP (1: 5000, Novus Biologicals, Cat. NB300-141, Centennial, Colorado, USA) or in a rat monoclonal antibody raised against microglia marker Iba1 (1:2000, Abcam, Cat. ab283346) for 18 hours at room temperature. Sections were next incubated in a biotinylated goat anti-mouse, in a biotinylated goat anti-rabbit IgG or in a biotinylated goat anti-rat IgG (1:200, MJS Biolynx Inc., Brockville, Ontario, Canada) for 1 hour and further processed by using an Elite ABC kit (MJS Biolynx Inc.) for another 1 hour according to the manufacturer instruction. Finally, the immunoprecipitates were developed using 3,3-diaminobenzidine (DAB, Sigma-Aldrich, Oakville, Ontario, Canada) as the chromogen and enhanced by the glucose oxidase-nickel-DAB method (Shu et al., 1988). Between incubations, PFC sections were washed thoroughly in PBS + T twice. Finally, sections were mounted on the pre-cleaned super plus glass slides (Fisher Scientific) or gelatin-coated slides, dehydrated in ascending ethanol solutions and cleared in xylene, and cover-slipped with xylene-based mounting medium (Micromount, Leica). After air drying overnight, sections were observed under the bright field of a microscope (Olympus Microscopes, 1X73, PIF, Montreal, Canada) and the digital images of immunoreactive cells were captured using the CellSens software (Olympus). Omission of primary antisera in all tested slides resulted in negative immunostaining.

### 2.4 Hemaetoxylin and eosin Y staining and Cresyl violet staining

On the day of HC staining, slides mounted with the 10-µm-thick PFC sections from the 5 groups were taken out from −80°C freezer and air dried for about 30 min to completely remove the moisture.

For H&E staining, sections were directly stained with 0.1% Mayers Hematoxylin solution (Sigma, Cat. MHS-16) for 10 min before bluing in warm running tape water (pH 7.8) for 5 min. Sections were stained with 0.5% ethanol eosin Y solution (Sigma, Cat. HT110116) for 30 sec. Sections were then dipped in ddH_2_O, 50% and 70% ethanol briefly, then dehydrated in 95% ethanol and 100% ethanol (2X, 5 min each) and cleared in xylene (2X, 5 min each). Finally, slides were cover-slipped, aired dried, and observed under the microscope as mentioned above. The nuclei of stained cells appeared as dark blue, the cytoplasm as pink while the erythrocytes were stained as bright red. The capillary and small blood vessels were stained as bright red which can be easily identified in the neuropils.

For CV staining, sections were incubated in 100% ethanol for 5 min and in 95% ethanol for 3 min. Then sections were rehydrated in 70% ethanol for 30 sec and in ddH_2_O for 30 sec. Sections were stained in 0.5% CV solution for 4 min. Sections were rinsed in ddH_2_O briefly before being differentiated in 70% acetified ethanol for 30 sec. Sections were then dehydrated in 90% and 100% ethanol (2X, 2 min each). Finally, sections were cleared in xylene (2X, 5 min each). Slides were cover-slipped, air dried and observed under the microscope as mentioned above. CV-stained Nissl substances in the cytoplasm of neurons and neuroglia appeared as purple.

### 2.5 Luxol fast blue-Eosin Y-Cresyl violet staining

On the day of LFB-EY-CV staining, 50µm-thick human PFC sections in cryoprotectant were taken out from −20°C freezer, rinsed in PBS, mounted on the gelatin-coated slides, and air dried for about 2 hours to completely remove the moisture. Sections were soaked in xylene (2X, 15 min each), 100% and 95% ethanol (2X, 5 min each), to de-fat the myelin in PFC sections. Sections were then incubated in 1% LFB solution (Luxol fast blue stain kit, Abcam, Cat. Ab150675) at 45°C overnight. On the second day, sections were briefly rinsed in 95% ethanol and ddH_2_O (5 sec each), differentiated in 0.05% lithium carbonate solution, 70% (2X, 30 sec each), and rinsed in ddH_2_O briefly. The differentiating procedures were repeated until a sharp contrast between the dark blue white matter and the colorless gray matter was observed. Briefly rinsed in 70% ethanol, sections were stained in 0.05% acidified ethanol EY for 30 sec. After a brief wash in ddH_2_O, sections were stained in 0.5% CV solution for 1 min. Then sections were rinsed in ddH_2_O briefly, dehydrated through 95% and 100% ethanol and cleared in xylene (2X, 5 min each). Section mounted slides were cover-slipped, air dried and observed under microscope as mentioned above. Myelin in the white matter of PFC sections were stained as dark blue or blue by LFB, but sparse or absent in the gray matter, particularly absent in the superficial layers. Neuropil or the cytoplasm of cells were stained as pink while the red blood cells in the blood vessels were stained as bright red by EY. The cytoplasm of neurons was specifically stained by CV as purple.

### 2.6 Image capture, quantification and statistical analysis for IHC- and HC-stained sections

In addition to the same staining procedures for all sections from the 5 groups, the setup for image capturing was consistent for all PFC sections from the 5 groups. Bright field images of IHC- or HC-stained profiles were captured at the same magnification (20X), with the same threshold of illumination or exposure time on the Olympus microscope (1X73 PIF). Each captured field results in an image with a dimension of 650µm x 475µm. The average optical intensity for each image was measured automatically using ImageJ (ImageJ ver.1.53, Softonic, Barcelona, Spain). The values of average optical intensity measured fall into the 1 to 256 range of the grayscale.

The intensity from the measurements from 5 images were averaged for each brain sample. The mean intensities from all groups (1, 5, 10, 15 and 20 years of post-fixation) were statistically compared by using one-way ANOVA with Student-Newman-Keuls multiple comparison tests using SigmaPlot V14 (Grafiti, St Palo Alto, California, USA). Linear regression analysis was performed to examine the relationships between the intensity of each staining and the post-fixation years, donor age and PMI for all samples from the 5 groups (n=20) using the software of SigmaPlot v14 (Grafiti). P and R values were determined by the software automatically. For all analyses, the significance level was set as p<0.05.

## 3. Results

### 3.1 Effect of prolonged post-fixation on NeuN, GFAP and Iba1 IHC staining

In the current study, we used two monoclonal antibodies raised against NeuN provided by Abcam (see Table 2). Using the mouse monoclonal anti-NeuN antibody (Msx) for the 50-µm-thick PFC section from the 5 post-fixation groups, we observed that abundant MsxNeuN-immunoreactive (IR) neurons were present in the gray matter of human brains post-fixed from 1 to 20 years (Fig 1A to 1E). MsxNeuN-IR neurons exhibited typical and clear neuron morphology with IR mainly localized in the nuclei, but also seen in the cytoplasm. From quantitative analysis, the MsxNeuN-IR intensity from the 10, 15 and 20-year-groups, but not from the 1-year-group, was significantly lower than from the 5-year-group (Fig. 1F, p>0.05). No significant difference was found among the 1, 10, 15 and 20-year-groups (Fig. 1F). Linear regression analysis showed a significant negative correlation between the intensity of MsxNeuN-IR neurons and the post-fixation years (Fig. 2A, p<0.03, R= −0.486).

**Figure 1.**
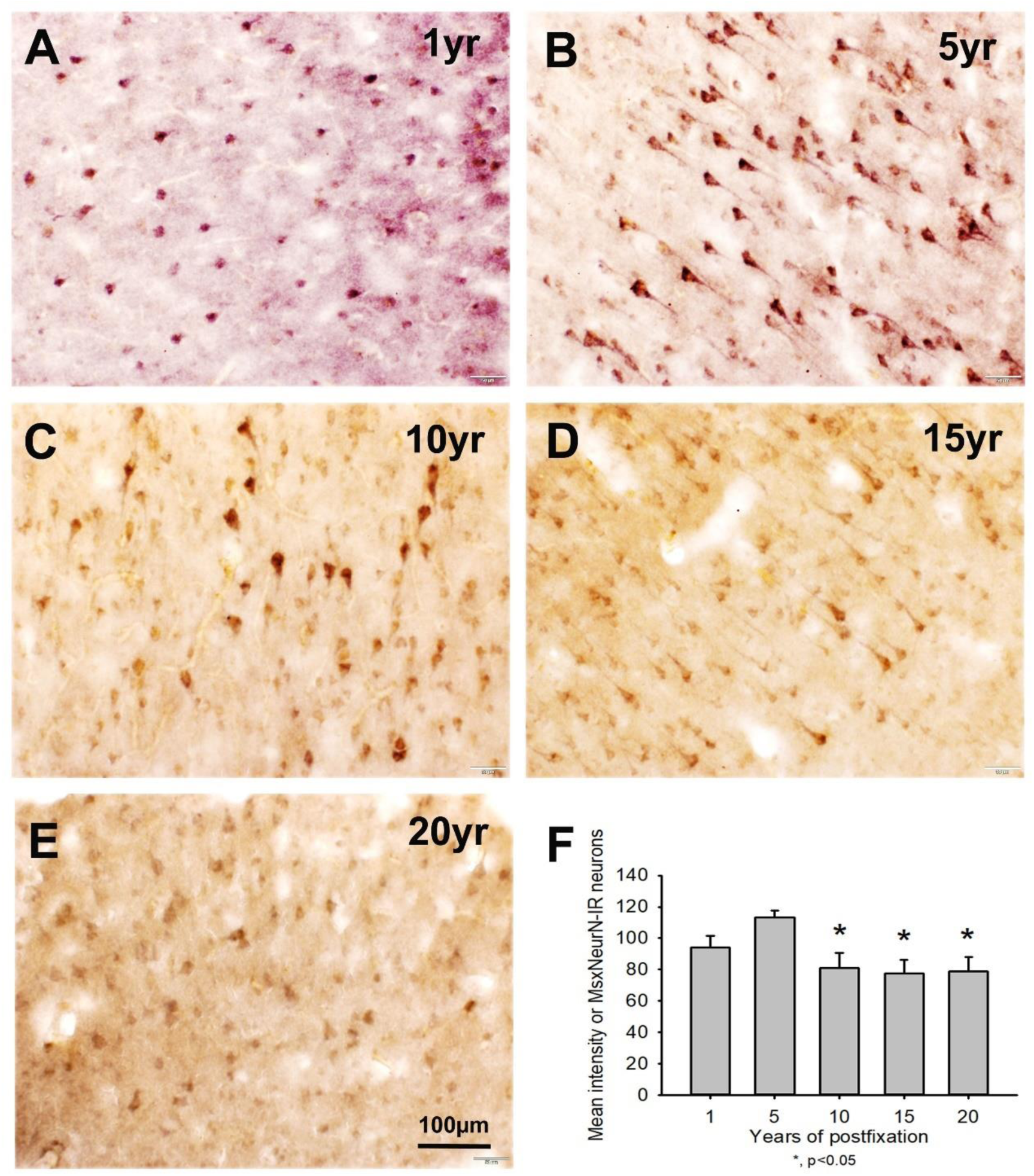
Expression of MsxNeuN-IR neurons in the gray matter of the 50-µm-thick PFC sections of human brains post-fixed for 1, 5, 10, 15 and 20 years. Numerous MsxNeuN-IR neuronal profiles were distributed throughout all layers in the gray matter of PFC sections of human brains post-fixed from 1 to 20 years (**A** to **E**). Immunoprecipitates were not only localized in the nuclei, but also in the cytoplasm of stained neurons. Quantitatively, the MsxNeuN-IR intensity from the 10, 15 and 20-year-groups, but not from the 1-year-group, was significantly lower than from the 5-year-group (**F**, * p<0.05). No significant difference was found among the 1, 10, 15 and 20-year-groups. Mean±SEM, n=4, one-way ANOVA with Student-Newman-Keuls multiple comparison test. The significance level was set at p<0.05.

**Figure 2.**
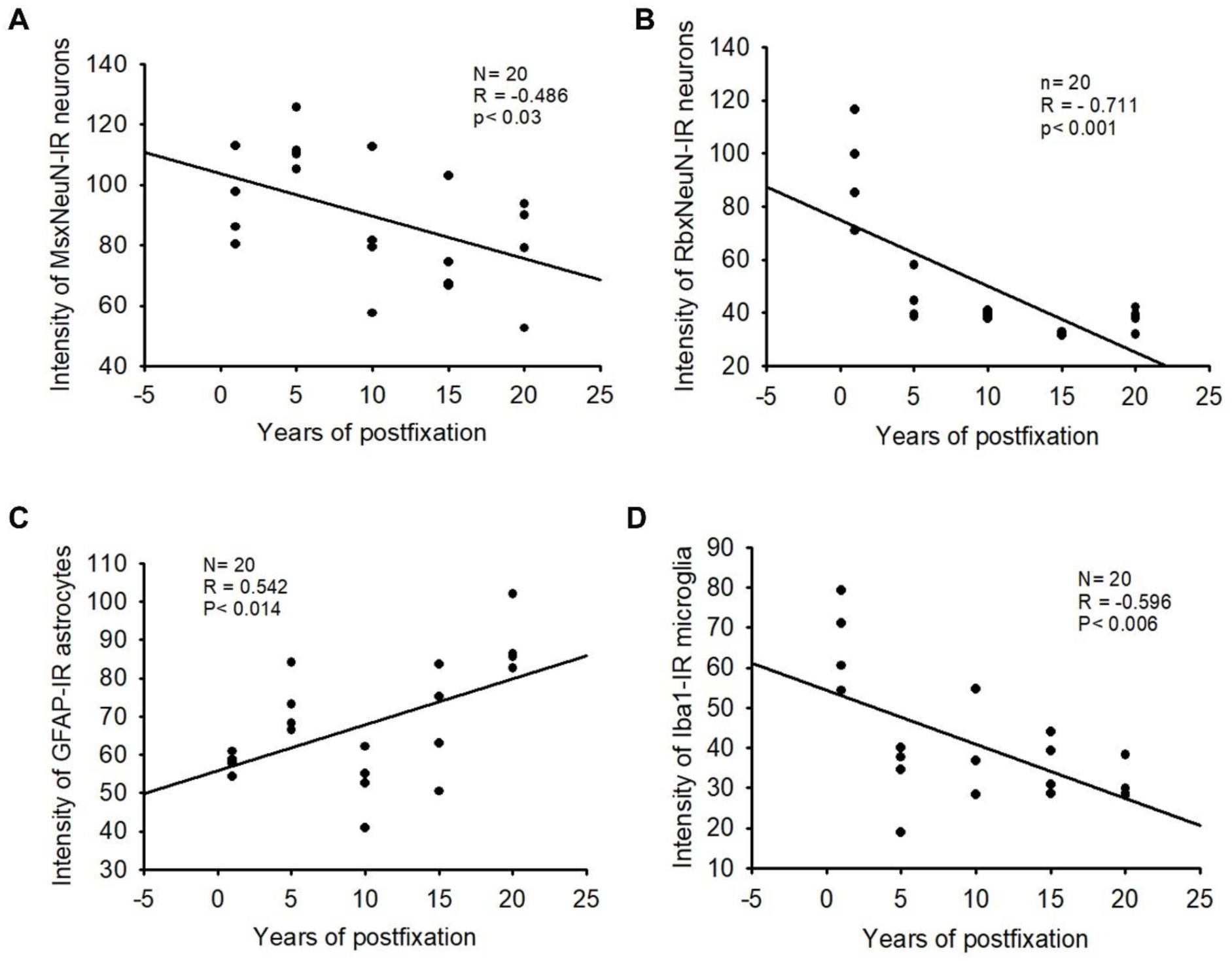
Linear regression analysis of the correlation between the intensity of MsxNeuN-IR neurons (**A)**, RbxNeuN-IR neurons (**B**), GFAP-IR astrocytes (**C**) and Iba1-IR microglia (**D**) in PFC of human brains verse the post-fixation years. A significant negative correlation was found between the intensity of MsxNeuN-IR neurons (**A**, p<0.03, R=-0.486), the intensity of RbxNeuN-IR neurons (**B**, p<0.001, R=-0.711), and the intensity of Iba1-IR microglia (**D**, p<0.006, R=-0.596) versus the post-fixation years. However, a significant positive correlation was detected between the intensity of GFAP-IR astrocytes and the post-fixation years (**C**, p<0.014, R=0.542). Linear regression analysis, n=20. Significance level was set as p< 0.05.

**Table 2.**
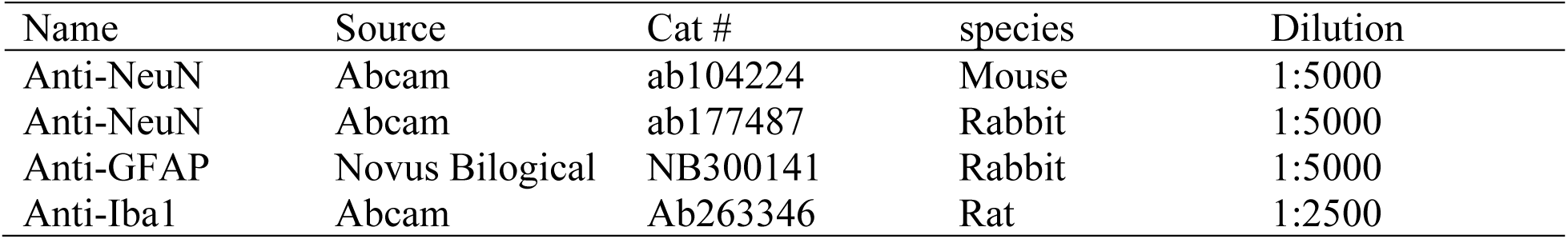
Summary of primary antisera used in this study.

| Name | Source | Cat # | species | Dilution |
| --- | --- | --- | --- | --- |
| Anti-NeuN | Abcam | ab104224 | Mouse | 1:5000 |
| Anti-NeuN | Abcam | ab177487 | Rabbit | 1:5000 |
| Anti-GFAP | Novus Biological | NB300141 | Rabbit | 1:5000 |
| Anti-Iba1 | Abcam | Ab263346 | Rat | 1:2500 |

We also used another rabbit monoclonal anti-NeuN antiserum (Rbx) from Abcam (Table 2) for the 50-µm-thick PFC section of the 5 groups. It was the same antibody used by a prior study showing a negative correlation of NeuN-IR and post-fixation time (Wu et al., 2022). Abundant RbxNeuN-IR neurons were only found in the gray matter of PFC sections from the 1-year-group (Fig 3A). Reliable RbxNeuN-IR neurons were invisible in the 5, 10, 15 and 20 year-groups (Fig 3B to 3E). Quantitatively, the intensity of RbxNeuN-IR neurons was significantly lower in PFC of the 5, 10, 15, and 20-year-groups than the 1-year-group (Fig. 3F, p<0.001). Linear regression analysis showed a significant negative correlation between the intensity of RbxNeuN-IR neurons and the post-fixation years (Fig. 2B, p<0.001, R=-0.711). Together, the above observations suggest that prolonged post-fixation reduces NeuN IHC staining for human brains.

**Figure 3.**
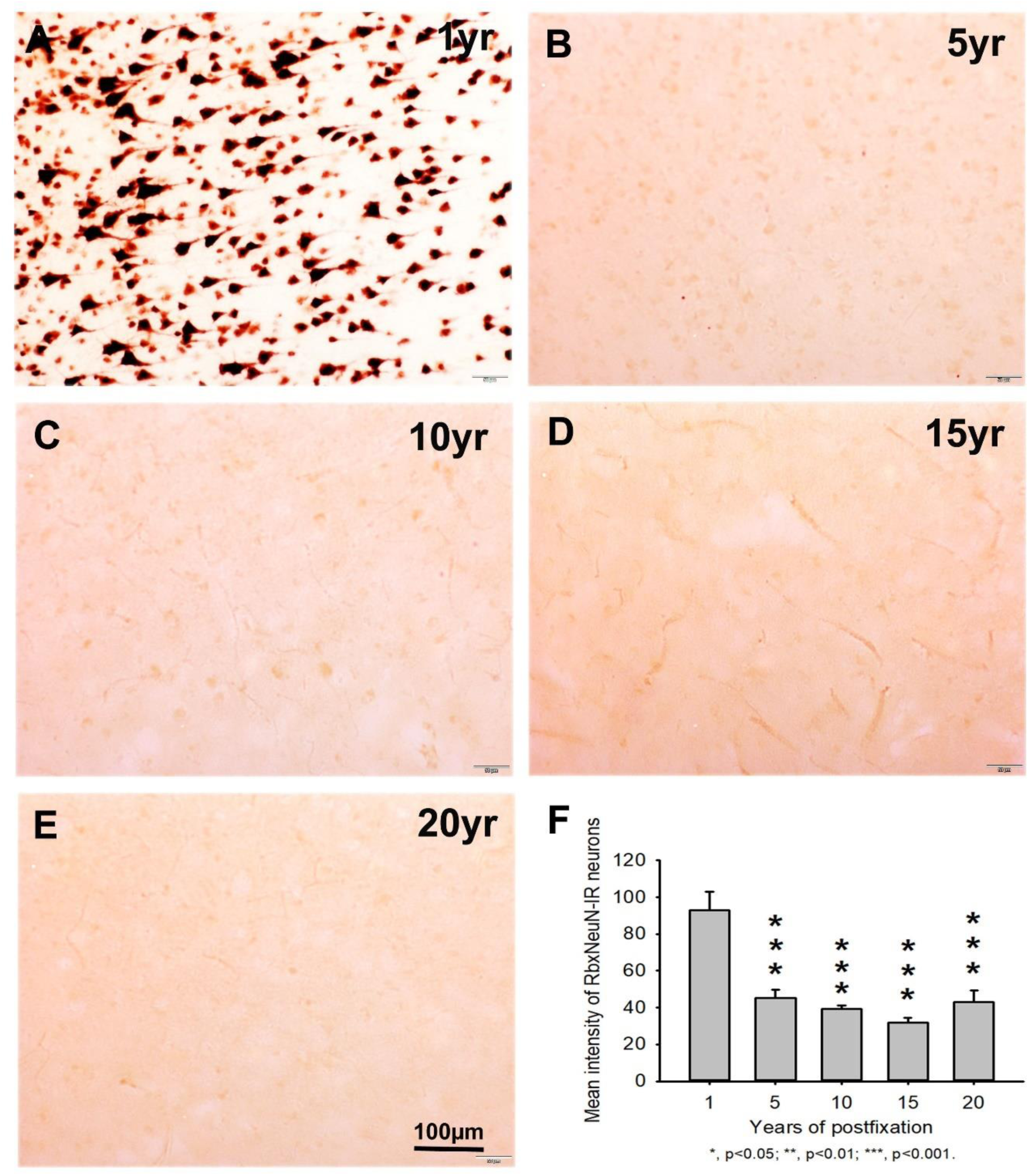
Expression of RbxNeuN-IR neurons in the gray matter of the 50-µm-thick PFC sections of human brains post-fixed for 1, 5, 10, 15 and 20 years. Abundant strongly stained RbxNeuN-IR neurons were only observed in PFC of the 1-year-group (**A**). However, reliable RbxNeuN-IR neurons were almost invisible in PFC of all other groups (**B** to **E**). Quantitatively, the intensity of RbxNeuN-IR neurons from the 5, 10, 15 and 20-year-groups was very significantly lower than from the 1-year-group (**F**, p<0.001). Mean±SEM, n=4, one-way ANOVA with Student-Newman-Keuls multiple comparison test. The significance level was set at p<0.05.

Using a rabbit anti-GFAP antibody from Novus Biological (Table 2), we found that GFAP-IR astrocytes were abundantly distributed throughout the gray and white matter of PFC section from human brains post-fixed for 1, 5, 10, 15 and 20 years (Fig 4A to 4E). GFAP-IR profiles exhibited typical characteristics of astrocytes, i.e., stellate soma with numerous rather long processes and branches. Quantitatively, the intensity of GFAP-IR astrocytes from the 1, 5, 10, 15-year-groups was significantly lower than from the 20-year-group (Fig. 4F, p<0.05 to <0.001). No significant difference was detected among the 1, 5, 10 and 15-year-groups (Fig.4F). Linear regression analysis disclosed a significant positive correlation between the intensity of GFAP-IR astrocytes and the post-fixation times (Fig 2C, p<0.014, R=0.542). These data suggest that prolonged post-fixation enhances GFAP IHC staining for human brains.

**Figure 4.**
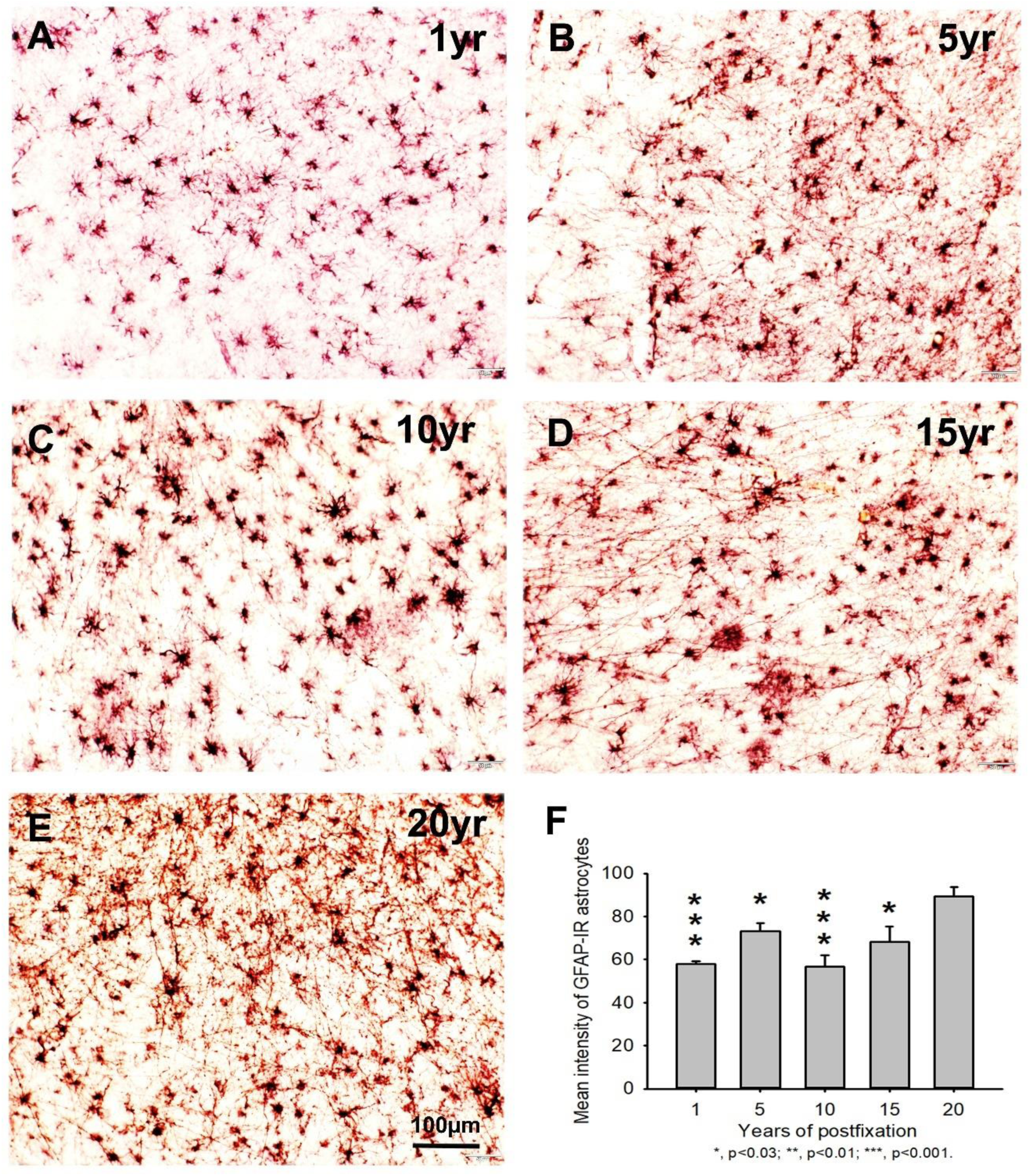
Distribution of GFAP-IR astrocytes in the 50-µm-thick PFC sections of human brains post-fixed for 1, 5, 10, 15 and 20 years. GFAP-IR astrocytes were abundantly distributed throughout the gray and white matter of PFC from human brain post-fixed for 1, 5, 10, 15 and 20 years (**A** to **E**). GFAP-IR astrocytes exhibited typical characteristics of astrocytes, a stellate soma with rather long and branched processes. Quantitative analysis revealed that the intensity of GFAP-IR astrocytes from the 1, 5, 10, 15-year-groups was significantly lower than from the 20-year-group (**F**, p<0.05 to 0.001). No significant difference was detected among the 1, 5, 10 and 15-year-groups (**F**). Mean±SEM, n=4, one-way ANOVA with Student-Newman-Keuls multiple comparison test. The significance level was set at p<0.05.

In the current study, we used a rat anti-Iba1 monoclonal antibody (Table 2) for IHC staining on the 50-µm-thick PFC sections of human brains post-fixed for 1, 5, 10, 15 and 20 years. We found that Iba1-IR microglia were observed throughout the gray and white matter of the PFC of human brain post-fixed for 1 to 20 years (Figs 5A to 5E). Iba1-IR microglia exhibited typical characteristics of microglia, with small irregular soma and multiple short processes. Quantitatively, the intensity of Iba1-IR microglia from the 5-, 10-, 15- and 20-year-groups was very significantly reduced compared to the 1-year-group (Fig 5F, p<0.001). No significant difference was detected among the 5-, 10-, 15- and 20-year-groups. Linear regression analysis revealed a significant negative correlation between the intensity of Iba1-IR microglia and the post-fixation years (Fig. 2D, p<0.006, R=-0.596). These observations suggest that prolonged post-fixation reduces Iba1 IHC staining for human brains.

**Figure 5.**
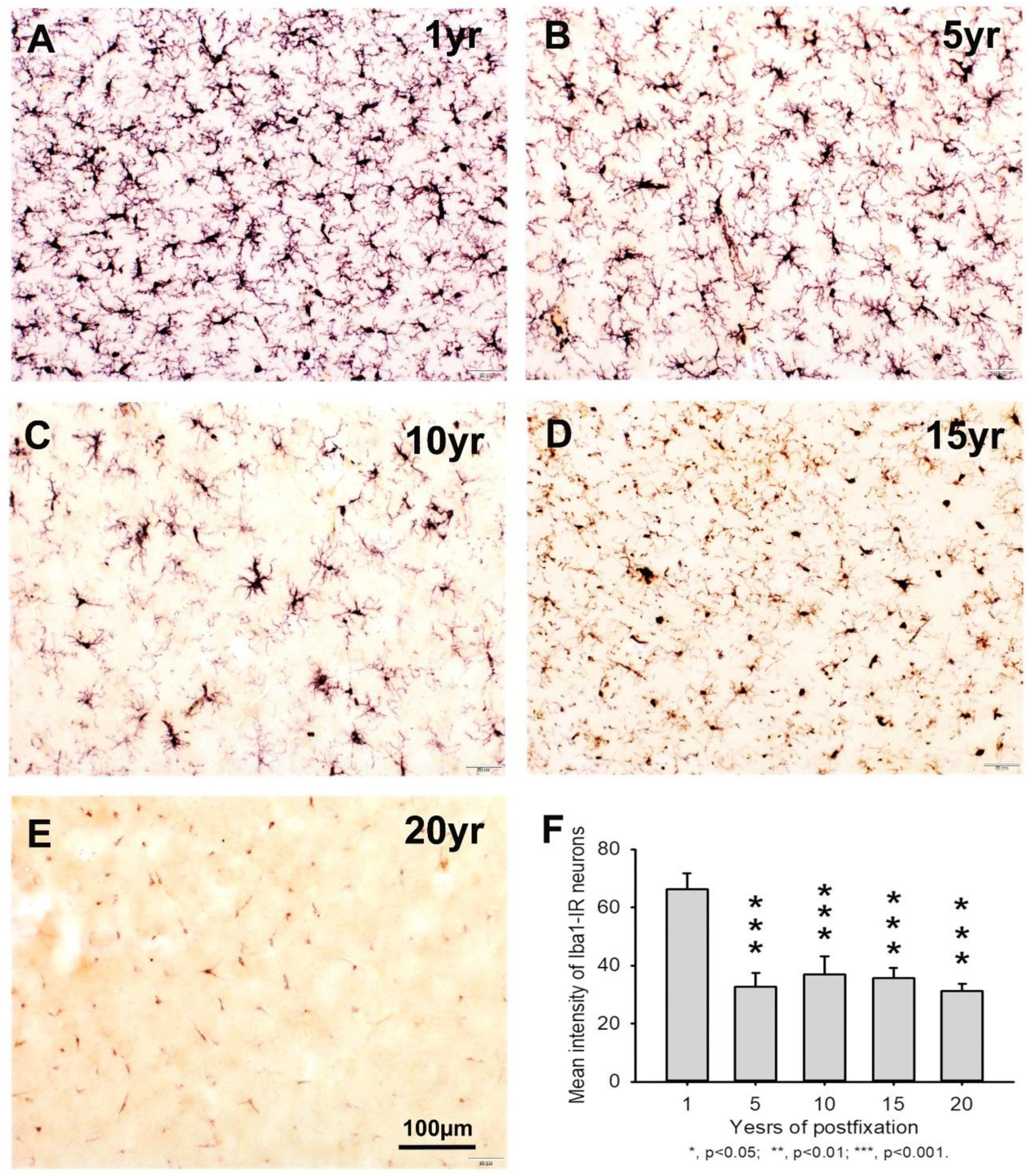
Distribution of Iba1-IR microglia in the 50-µm-thick PFC sections of human brains post-fixed for 1, 5, 10, 15 and 20 years. Iba1-IR microglia were distributed throughout the gray and white matter of PFC of human brain post-fixed for 1 (**A**), 5 (**B**), 10 (**C**), 15 (**D**) and 20 (**E**) years. Quantitatively, the intensity of Iba1-IR microglia from the 5-, 10-, 15- and 20-year-groups was very significantly reduced compared to that from the 1-year-group (**F**, p<0.001). No significant difference was detected among the 5-, 10-, 15- and 20-year-groups. Mean±SEM, n=4, one-way ANOVA with Student-Newman-Keuls multiple comparison test. The significance level was set at p<0.05.

### 3.2 Effect of prolonged post-fixation on hemaetoxylin & eosin Y staining

In this study, we examined H&E staining for the 10-µm-thick PFC sections of human brains post-fixed for 1, 5, 10, 15 and 20 years. The nuclei of H&E-stained cells were dark blue while the cytoplasm was pink (Fig. 6A to 6E). In contrast, red blood cells in the small vessels (Fig 6A to 6E, arrows) and capillaries (Fig 6D and 6E, arrowheads) were stained as bright red in the neuropils. Quantitatively, the intensity of H&E staining from the 5- and 15-year-groups was significantly higher than from the 1-year-group (Fig. 6F, p<0.05 and p<0.001, respectively). No significant difference was detected between any other two groups (Fig. 6F). Linear regression analysis showed a significant positive correlation between the intensity of H&E staining and the post-fixation years (Fig. 7A, p<0.011, R=0.553). These data suggest that prolonged post-fixation enhances H&E staining for human brains.

**Figure 6.**
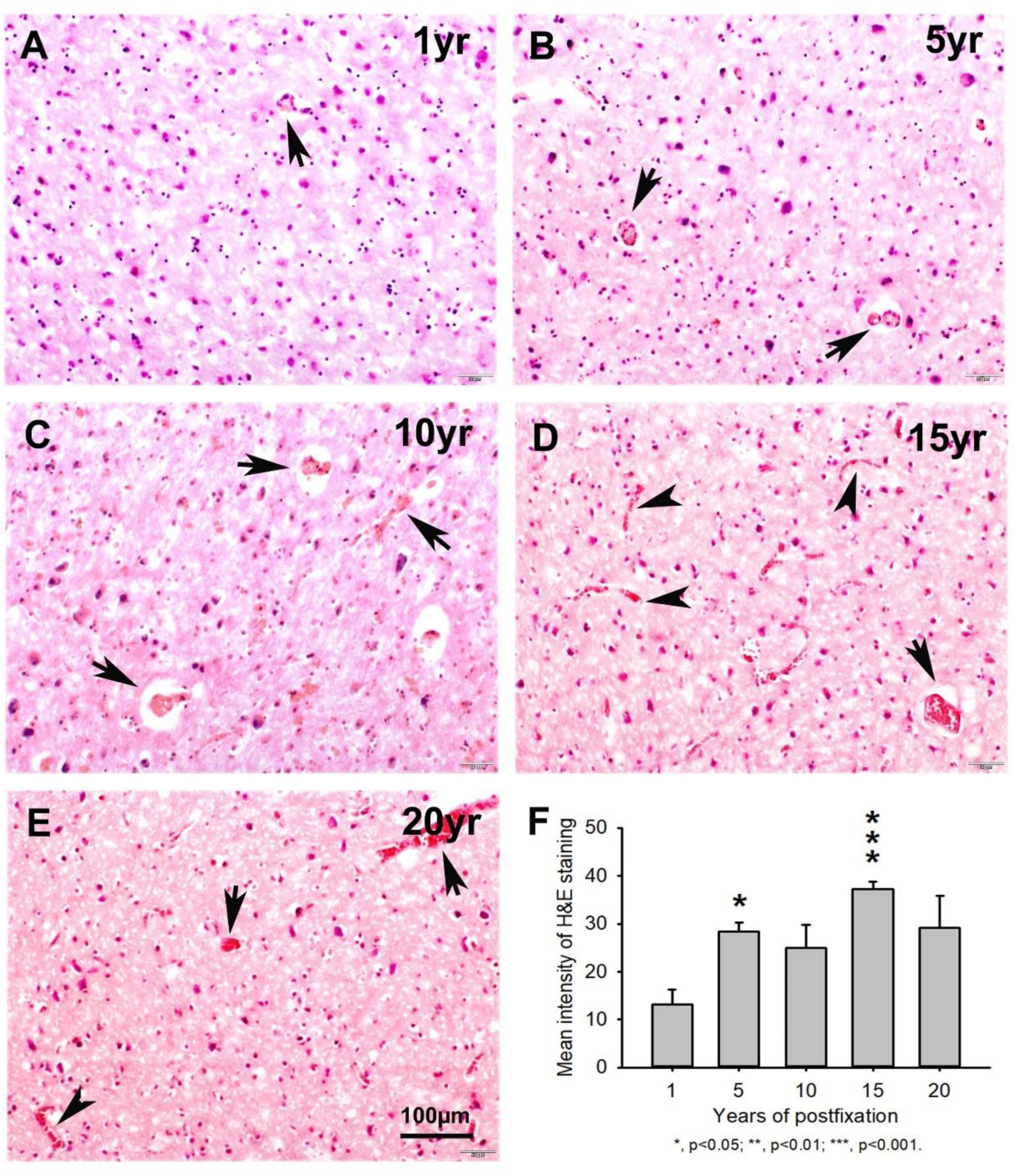
Hemaetoxylin (H) and Eosin Y (E) staining for the 10-µm-thick PFC sections of human brains post-fixed for 1, 5, 10, 15 and 20 years. In H&E-stained PFC sections (**A** to **E**), the nuclei of cells were stained as dark blue by H while the cytoplasm was stained as pink by E. In contrast, the red blood cells in the small vessels (**A** to **E**, arrows) and capillaries (**D** and **E**, arrowheads) were stained by E as bright red, which is easy to identify in the neuropils. Quantitatively, the intensity of H&E staining from the 5- and 15-year-groups was significantly higher than from the 1-year-group (**F,** p<0.05 to 0.001). There was no significant difference between any other two groups (**F**). Mean±SEM, n=4, one-way ANOVA with Student-Newman-Keuls multiple comparison test. The significance level was set at p<0.05.

**Figure 7.**
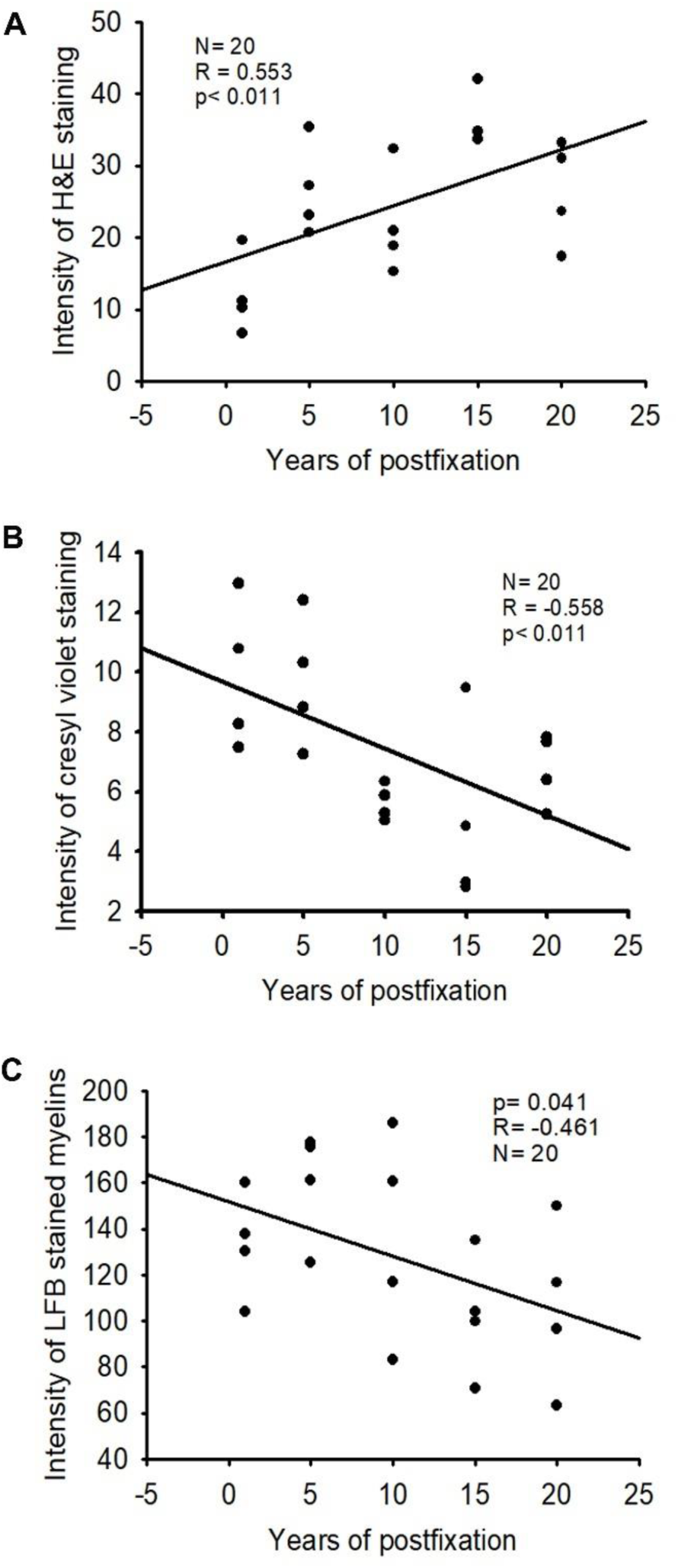
Linear regression analysis of the correlation between the intensity of H&E staining (**A**), CV staining (**B**) or LFB staining (**C**) in PFC of human brains and the post-fixation years. A significant positive correlation was detected between the intensity of H&E staining and the post-fixation years (**A,** p<0.011, R= 0.553). A significant negative correlation was observed between the intensity of CV staining (**B,** p<0.011, R= 0.558) or LFB staining (**C,** p<0.041, R= −0.461) and the post-fixation years. Linear regression analysis, n=20. Significance level was set as p< 0.05.

### 3.3 Effect of prolonged post-fixation on cresyl violet staining

We performed CV staining for the 10-µm-thick PFC sections from human brains post-fixed for 1, 5, 10, 15 and 20 years. CV-stained neurons were observed in the gray matter of PFC sections from all groups (Fig. 8A to 8E). Nissl substances in the cytoplasm of neurons was stained as purple or dark purple. The abundance and intensity of CV-stained neurons were detected among the 1- and 5-year-groups (Fig 8A and 8B). However, the abundance and intensity of CV-stained neurons in PFC of the 10-, 15- and 20-year-group were evidently reduced (Fig 8C to 8E). Quantitatively, the intensity of CV staining from the 10, 15 and 20-year-groups was significantly lower than from the 1 (Fig. 8F, p<0.05) and 5-year-groups (Fig. 8F, p<0.05). No significant difference was detected among the 10, 15 and 20-year-groups. Linear regression analysis revealed a very significant negative correlation between the intensity of CV staining and the post-fixation years (Fig. 7C, p<0.011, R=-0.558). These data suggest that prolonged post-fixation reduces CV staining for human brains.

**Figure 8.**
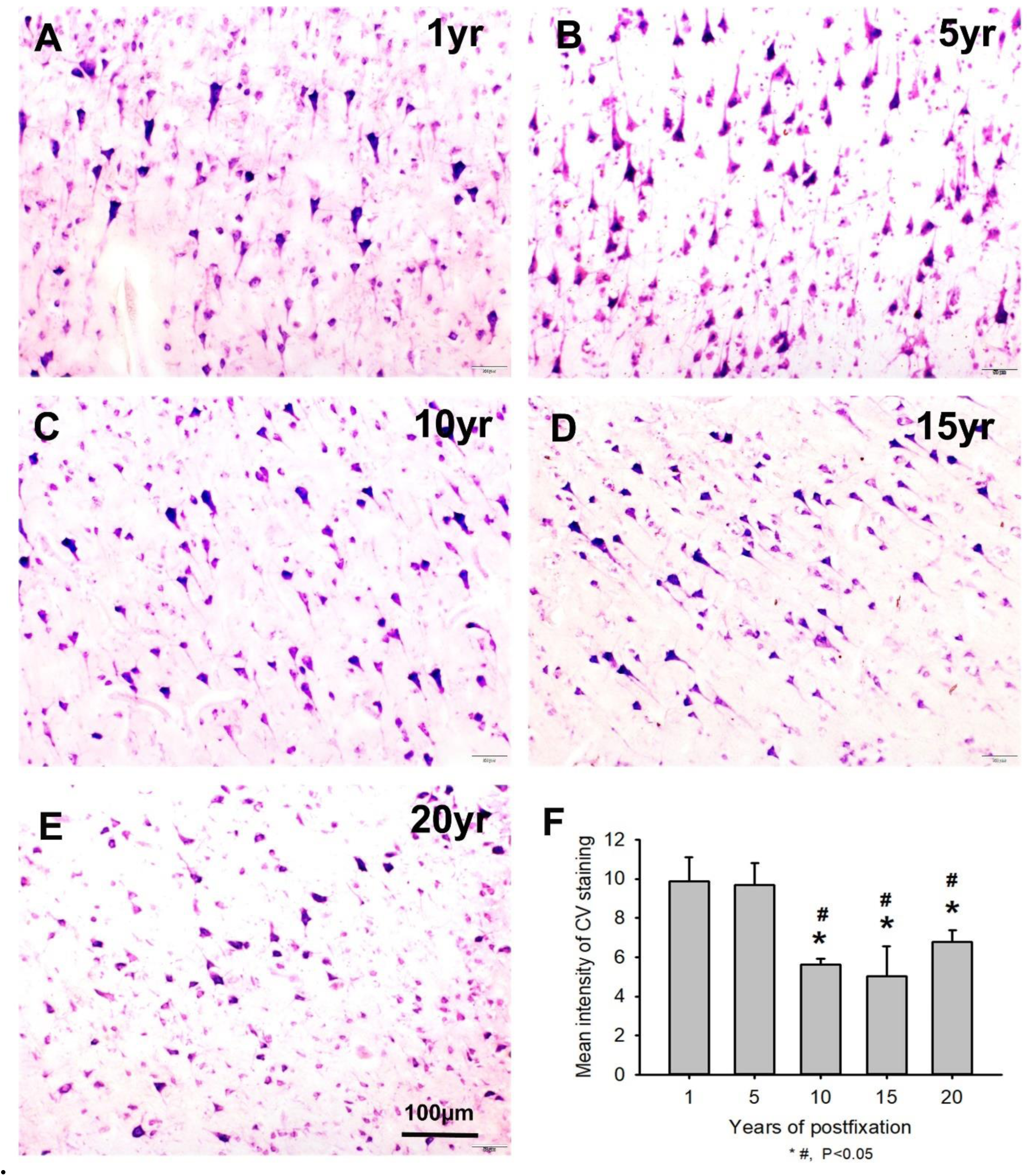
Cresyl violet staining of the 10µm-thick PFC sections of human brain post-fixed for 1, 5, 10, 15 and 20 years. In CV-stained PFC sections (**A** to **E**), the cytoplasm of neurons was stained as purple. Quantitatively, the intensity of CV staining from the 10-, 15- and 20-year-groups was significantly lower than from the 1- (**F**, *, p<0.05) and 5-year-groups (**F**, #, p<0.05). No significant difference was detected among the 10-, 15- and 20-year-groups. Mean±SEM, n=4, one-way ANOVA with Student-Newman-Keuls multiple comparison test. The significance level was set at p<0.05.

### 3.4 Effect of prolonged post-fixation on myelin staining

In this study, using LFB-EY-CV staining protocol, we performed myelin HC staining for the 50-µm-thick PFC sections from human brains post-fixed for 1, 5, 10, 15 and 20 years. We observed that the myelin enriched white matter of PFC from all brains of the 5 groups were stained as dark blue or blue by LFB while the neuropils and blood vessels were stained as pink and red by EY (Fig 9A to 9E). In the gray matter (data not shown), where myelin was absent or sparse, neurons were stained by CV as purple. Quantitatively, the myelin intensities from the 15- and 20-year groups appeared to be lower than those from the 1-, 5- and 10-year groups, but these differences were not significant on statistical analysis (Fig. 9F). However, linear regression analysis revealed a significant negative correlation between the myelin intensity and the post-fixation years (Fig. 7C, R= −0.461, p<0.041,). These data suggest that prolonged post-fixation reduces myelin staining of human brains.

**Figure 9.**
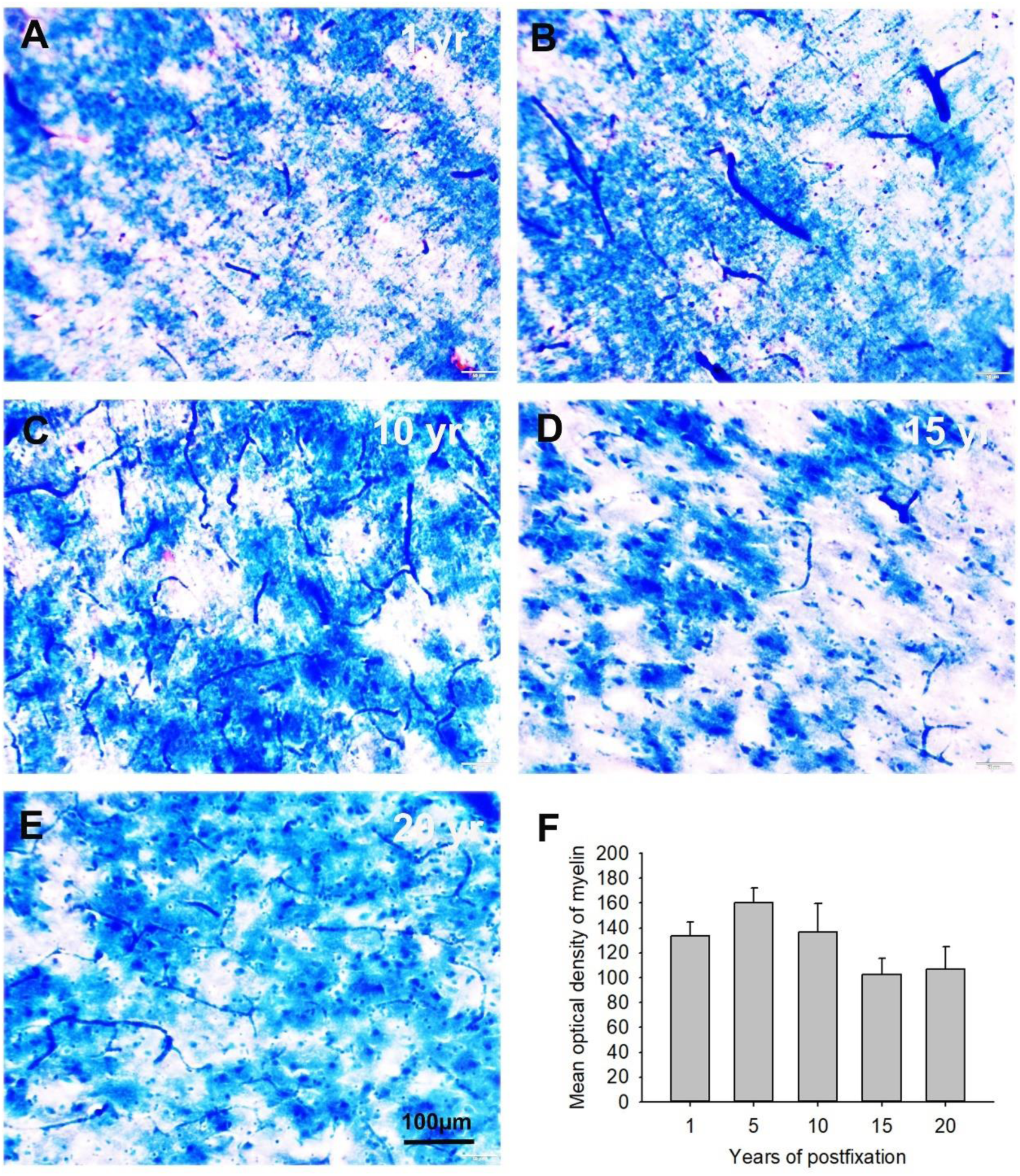
Luxol fast blue (LFB)-Eosin Y(EY)-cresyl violet (CV) staining of the 50µm-thick PFC sections from human brain post-fixed for 1, 5, 10, 15 and 20 years. In the myelin enriched white matter of PFC (**A** to **E**), heavily LFB-stained myelin profiles were observed. The neuropil and blood vessels were counterstained by EY as pink and bright red, respectively (**A** and **C**). Neurons in the myelin lacking gray matter was stained as purple (not shown here). Quantitatively, the myelin intensity in PFC from the 15 and 20-year groups were lower than those from the 1-, 5- and 10-year groups, but these differences were not statistically significant (**F**). Mean±SEM, n=4, one-way ANOVA with Student-Newman-Keuls multiple comparison test. The significance level was set at p<0.05.

### 3.5 Effects of donor age and postmortem interval on IHC and HC staining

In the current study, to examine the possible effect of donor age and PMI of brain samples on IHC and HC staining we performed linear regression analysis (Tables 3 and 4). There was a significant negative correlation between the donor age and the intensity of GFAP-IR astrocytes in PFC of human brains (Table 3, R=-0.483, p<0.031). However, there was no significant correlation between the donor age and the intensity of any other IHC or HC staining (Table 3). Moreover, we did not observe any significant correlation between PMI and the intensity of any IHC and HC staining (Table 4). These data suggest that donor age and PMI exert limited effect on IHC and HC staining.

**Table 3.** Summary of linear regression analysis between ages and intensity of IHC and HC.

| Intensity | Age | Size | R | P |
| --- | --- | --- | --- | --- |
| MsxNeuN-IR | 19-91 | 20 | 0.0914 | 0.702 |
| RbxNeuN-IR | 19-91 | 20 | 0.390 | 0.089 |
| GFAP-IR | 19-91 | 20 | -0.483 | 0.031 |
| Iba1-IR | 19-91 | 20 | 0.179 | 0.449 |
| H&E staining | 19-91 | 20 | 0.139 | 0.559 |
| CV staining | 19-91 | 20 | 0.269 | 0.251 |
| LFB staining | 19-91 | 20 | 0.198 | 0.403 |

Footnotes: A significant negative correlation was detected between GFAP-IR intensity and age (R=-0.483, p<0.031). There was no significant correlation between intensity of any other IHC and HC staining versus age.

**Table 4.** Summary of linear regression analysis between PMI and intensity of IHC and HC.

| Intensity | PMI (hours) | Size | R | P |
| --- | --- | --- | --- | --- |
| MsxNeuN-IR | 10.72 - 98.75 | 20 | 0.0678 | 0.777 |
| RbxNeuN-IR | 10.72 - 98.75 | 20 | 0.217 | 0.358 |
| GFAP-IR | 10.72 - 98.75 | 20 | 0.257 | 0.275 |
| Iba1-IR | 10.72 - 98.75 | 20 | 0.153 | 0.519 |
| H&E staining | 10.72 - 98.75 | 20 | 0.193 | 0.416 |
| CV staining | 10.72 - 98.75 | 20 | 0.0583 | 0.807 |
| LFB staining | 4.49 - 98.75 | 20 | 0.0630 | 0.792 |

Footnotes: There was no significant correlation between intensity of any IHC and HC staining versus PMI.

## 4. Discussion

### 4.1 Summary of major findings

In the current study, we examined NeuN, GFAP and Iba1 IHC staining as well as H&E, CV and LFB HC staining on PFC sections from human brains post-fixed for 1, 5, 10, 15 and 20 years. A negative correlation was detected between the IHC staining of NeuN and Iba1and the HC staining of CV and LFB versus the post-fixation years. By contrast, a positive correlation was found between GFAP IHC staining and H&E HC staining versus the post-fixation times. These data suggest that prolonged post-fixation exerts differential facilitating and inhibiting effects on IHC and HC staining. Furthermore, except for a negative correlation between the donor age and GFAP IHC staining, no significant correlation was detected between donor age or PMI versus any IHC and HC staining mentioned above. These data suggest that unlike post-fixation durations, donor age and PMI play limited roles in IHC and HC staining for postmortem human brains.

### 4.2 Effects of prolonged post-fixation on NeuN, GFAP and Iba1 IHC staining for human brains

Prior studies showed a negative correlation between NeuN-IR intensity and post-fixation times ranging from immediate to 3 years (Wu et al., 2022) or from post-fixation durations of 24 hours, 4 months and 10 years (Lyck et al., 2008). Another prior IHC study on post-fixed swine brain showed that NeuN expression was reduced in swine brain post-fixed for 2 months, but completely disappeared after 3-month-post-fixation (Lundstrom et al., 2019). These prior findings suggest that prolonged post-fixation even within a couple of months or years suppresses NeuN expression in IHC staining. In this study, we greatly expanded the time span of examination for human brain tissues. Consistent with these prior studies, we detected a negative correlation between NeuN-IR intensity and post-fixation time on human brains by using two anti-NeuN antisera from Abcam. However, the two antisera resulted in rather different IHC staining patterns although the trend of post-fixation exerted effect was similar. Reliable NeuN-IR neurons stained with a mouse monoclonal antibody appeared in all groups, even visible in brains from the 20-year-group. However, NeuN-IR neurons stained with a rabbit monoclonal antibody, the same antibody used in the study by (Wu et al., 2022), only existed in brains from the 1-year-group. Our data suggest that the effect of prolonged post-fixation might also depend on used antibodies or relevant molecular changes in tissues. A suitable NeuN antibody could reveal target protein even in brain samples post-fixed for two decades although the staining intensity was significantly reduced compared to samples from early years of post-fixation. As such, it is possible to compensate for the prolonged post-fixation reduced immunoreactivity with a more effective antibody. It is recommended to assess more available antibodies and find the most optimal antibody to perform IHC staining for postmortem brains, thus offsetting the negative effect of prolonged post-fixation on IHC staining.

GFAP is an important biomarker widely used in diagnostic and research of neurological disorders such as Alzheimer disease (Abdelhak et al., 2022; Kim et al., 2023). Astrocytosis could be an alternative pathway for the pathogenesis of AD (Pereira et al., 2021). To understand the mechanisms underlying neurological disorders, it is important to examine GFAP expression in archived human AD brain post-fixed for years and decades. Previously, it was shown that GFAP levels was reduced in human brains with post-fixation ranging from 24 hours to 4 months and up to 10 years by immunoarray, but GFAP-IR astrocytes were also sufficiently seen in tissue fixed for 10 years when applying HIER in IHC (Lyck et al., 2008). Another study showed that GFAP-IR intensity remained unaltered in human brains post-fixed ranging for months up to 3 years by IHC staining (Wu et al., 2022). Contrary to these studies, for the first time we observed here a positive correlation between GFAP-IR intensity and post-fixation time ranging over 1, 5, 10, 15 and 20 years. We further observed that the mean GFAP-IR intensity from the 20-year-group was significantly higher than other 4 groups, but no significant difference was detected among other 4 groups. Indeed, we also observed abundant strongly stained GFAP-IR astrocytes in PFC and hippocampus of human brain post-fixed for 25 years (n=1, supplemental Fig. 1). Our data strongly suggest that prolonged post-fixation exerts a positive or facilitating effect on GFAP IHC staining. Detection of GFAP in human brains post-fixed for >20 years definitively paves a novel way for the use of large quantities of donated and archived AD brains to uncover the underlying mechanisms of this deteriorating disease.

Elevation of microglia biomarker is a consistent feature of AD brains (Hopperton et al., 2018), suggesting a crucial role of microglia activation in AD pathogenesis. Thus it is important to determine the effect of post-fixation on microglia marker such as Iba1 in IHC studies. Prior studies using immunoassay showed that active microglia marker (CD45+ and CD68) in postmortem human brains was negatively correlated with post-fixation time (from 24hr to 4 months and up to 10 years) (Lyck et al., 2008). However, a subsequent study showed no alteration in Iba1 IHC staining detected in human brains post-fixed from weeks up to 3 years (Wu et al., 2022). Consistent with the study by Lyck et al (Lyck et al., 2008), in the current study, we found a very significant negative correlation between Iba1-IR intensity and post-fixation years. The mean intensity of Iba1-IR microglia was significantly reduced in human brains post-fixed for 5, 10, 15 and 20 years compared to the 1-year-group. These data suggest that extended post-fixation duration reduces Iba1 IHC expression of human brains.

It remains unknown why prolonged formalin post-fixation exerts differential positive and negative effects on IHC staining. For the negative effect, it was assumed that prolonged post-fixation progressively enhances the crosslinking between proteins to mask most antigen epitopes, thus leading to the reduced immunoreactivity. On the contrary, formalin induced protein crosslinking in human tissues might change protein conformation to expose some antigens. However, in our pioneer study (Supplemental Fig. 1), we observed that citrate buffer (pH6.0) based HIAR, which breaks up formalin induced crosslinks, enhanced GFAP IHC staining for brains post-fixed for 25 years. This observation indicates that formalin induced crosslinks is unlikely to be the reason for the positive effect of prolonged post-fixation on GFAP IHC staining. Moreover, we still observed that prolonged post-fixation also exerts differential positive and negative effects on HC staining (see below), which further support the assumption that more antigen exposures by formalin induced crosslinks is unlikely the reason for the positive effect.

Prior studies suggest that molecular-dependent changes occurring in prolonged formalin-fixed brain tissues are likely associated with the effect of post-fixation on IHC (Lyck et al., 2008; Wu et al., 2022). Indeed, it was shown that the effects of post-fixation delays >12 hour specifically relied on each antigen since IR-cells could be artificially increased, decreased, or unaffected depending upon the targeted antigen (Lyck et al., 2008; Engel and Moore, 2011). Based on these findings, the differential positive and negative effect of prolonged post-fixation on IHC staining might be the reflection of the molecular changes due to formalin or other factors in brain tissues.

### 4.3 Effects of prolonged formalin post-fixation on H&E, CV and LFB HC staining for postmortem human brains

Hemaetoxylin and eosin (H&E) staining is the standard histological protocol as it produces a good overview of the tissue and the cellular components to clearly display different types of structures. In the current study, for the first time we demonstrated a positive correlation between H&E staining intensity and post-fixation time of 1, 5, 10, 15 and 20 years. This observation suggest that post-fixation improves H&E HC staining.

Cresyl violet (CV) staining is a fundamental method to visualize the rough endoplasmic reticulum and ribosomes, the Nissl substance, along the course of dendrites or in neuronal perikaryons. Regarding the effect of prolonged post-fixation on CV staining, it was previously shown that intensity of CV-stained neurons was negatively correlated with post-fixation time (from weeks to 3 years) (Wu et al., 2022). In the current study, we greatly expanded the time span to examine this issue and consistently found a negative correlation between CV staining intensity and post-fixation durations ranging over 1, 5, 10, 15 and 20 years.

In addition to magnetic resonance imaging (MRI) and IHC staining for myelin basic protein (MBP), Luxol fast blue (LFB) staining is a most common HC technique to examine myelin and axonal degeneration in diagnostic and research. Prior studies showed that formalin post-fixation of postmortem human brains for 43 and 64 days altered MRI indices of myelin images (Schmierer et al., 2010; Shatil et al., 2018). Using tissue immunoassay, it was reported that MBP-IR was reduced in human brains post-fixed from 4 months to 10 years (Lyck et al., 2008). Another more recent qualitative study showed that prolonged post-fixation of human brains (a group of 1 to 20 years) reduced myelin intensity on LFB staining compared to the control group (< 1 year) (Alrafiah and Alshali, 2019). In this study, we addressed this issue in a more systematic and quantitative way and found a negative correlation between myelin intensity and post-fixation time (1, 5, 10, 15, and 20 years) on regression analysis. However, using one-way ANOVA with multiple comparison test, no significant difference in the myelin intensity was detected among the 5 groups, although myelin intensity of the 15- and 20-year-groups were evidently lower than these of the 1, 5 and 10 year-groups. These data suggest that prolonged post-fixation reduces myelin staining, but this reduction in the first 10 years was not evident.

Taking above mentioned data into consideration, a question was raised as to why prolonged post-fixation exerts differential positive and negative effects on HC staining. Formalin induced protein crosslinks seem suitable to explain a stronger H&E staining as post-fixation prolongs. But this presumption fails to interpret the negative effect of prolonged post-fixation on CV and LFB staining. Alternative theories must be explored such as degradation of certain proteins such as myelin or structures as well as Nissl substances. But the loss of NeuN stained neurons and Iba1 stained microglia is surpassed by the gain of GFAP stained astrocytes or H&E-stained other cell types so that an enhanced H&E staining could be noticed. Further studies are needed to address these issues.

It is worth mentioning that most prior studies exploring the effect of prolonged post-fixation on IHC and HC staining were performed on paraffin-embedded human brain sections. The current study for the first time examined this issue on both thin and thick cryosections of postmortem human brains. Although paraffin embedding provides better tissue preservation, its process also prevents the penetration of antibodies (Ramos-Vara et al., 2014). Cryosectioning is a fast and convenient technique to obtain large quantities of thin (10 to 20 µm thick) and thick (40 to 50µm) sections in a short time. IHC for mounted thin sections or for free-floating thick sections makes staining more versatile and flexible. Following HIAR, the quality of IHC staining was comparable for both paraffin embedded sections and cryosections (Shi et al., 2011). It was also shown that formalin post-fixation and subsequent cryosectioning did not affect the size and shape of ocular tissues (Tran et al., 2017). Our data support the reliability of using cryosections from human brains post-fixed for years and decades in research of neurological and psychiatric disorders.

### 4.4 Effects of donor age and postmortem interval on IHC and HC staining

In this study, except for a negative correlation between age and GFAP IHC staining, no significant correlation between age or PMI versus IHC and HC staining was observed. Our data were consistent with previous studies showing no correlation between PMI and IHC. For example, it was shown that immunostaining profiles for most proteins remained unchanged even after PMI of over 50 hours (Blair et al., 2016) and cellular features appeared to remain intact for more extended PMI (Krassner et al., 2023). Even when RNA was degraded, the protein levels in postmortem human remained stable (Stan et al., 2006). Furthermore, we recently showed that PMI had no significant effects on histology quality (Frigon et al., 2024). All these data suggest that PMI up to 3 days likely exerts no major effects on IHC and HC staining of postmortem brains. A recent review mentioned that the evidence for the interaction of age with PMI effects is mixed and may depend on the structural feature considered (Krassner et al., 2023). Isolation of primary microglia from the human post-mortem brain was not affected by donor age and PMI (Mizee et al., 2017). However, a negative correlation was reported between age and immunoreactivity for G-protein subunit G_αi1/2_-, but not for subunit G_αi3_-, G_αo_-, and G_αs_-proteins (Gonzalez-Maeso et al., 2002). In this study, we only detect a negative correlation between age and GFAP-IR, but not other IHC and HC staining. GFAP expression was known to be increased in aging human brains (Wruck and Adjaye, 2020; Chatterjee et al., 2021). Elevated GFAP expression is likely involved in the pathogenesis of neurodegenerative diseases such as Alzheimer’s disease, Parkinson’s disease and amyotrophic lateral sclerosis (Palmer and Ousman, 2018). However, in our brain samples without any neurological disorders, we found that GFAP IHC staining was negatively affected by donor age, but positively affected by post-fixation years. Hence, to achieve consistent results, it is recommended to perform GFAP IHC staining on postmortem brain within similar ranges of donor age and post-fixation durations.

### 4.5 Significance of the findings from the current study

We demonstrated here a positive effect of prolonged post-fixation on GFAP IHC staining and H&E HC staining as well as a negative effect on NeuN, Iba1 IHC staining and CV and IFB HC staining for PFC from postmortem human brains. The differential positive and negative effects could result from the molecular conformational changes associated with post-fixation times. Moreover, we found no correlation between the donor age or PMI with IHC and HC staining for most target proteins or structures, except for GFAP. Thus, successful IHC and HC staining depends on post-fixation times, target molecules, available antibodies and staining procedures. Therefore, it is recommended to perform IHC and HC staining for postmortem human brains post-fixed at the same time windows, to use the most optimal antibodies and staining procedures. Since prolonged post-fixation exerts both positive and negative effects on IHC and HC staining, depending on the target molecules, it is necessary or unnecessary, depending on the examined targets, to perform these procedures as soon as possible after post-fixation is initiated. Since all IHC and HC staining was performed on thin or thick cryosections and reliable staining results were achieved, this study also supports the application of cryosectioning in morphological studies of postmortem brains.

The current study expands the time span of possible application of archived human brains post-fixed for years and decades for clinical diagnosis and brain research. Hence the data collected from the current study could be used to model and adjust for the impact of post-fixation on some of IHC and HC staining measures, if not all. Exploration of these long archived and post-fixed human brains will advance our understanding the mechanisms underlying neurological and psychiatric disorders.

## Acknowledgements

The PDF of this manuscript has been available on biorxiv website: https://www.biorxiv.org/content/10.1101/2024.05.13.593904v1

---

**Supplemental Figure 1.**
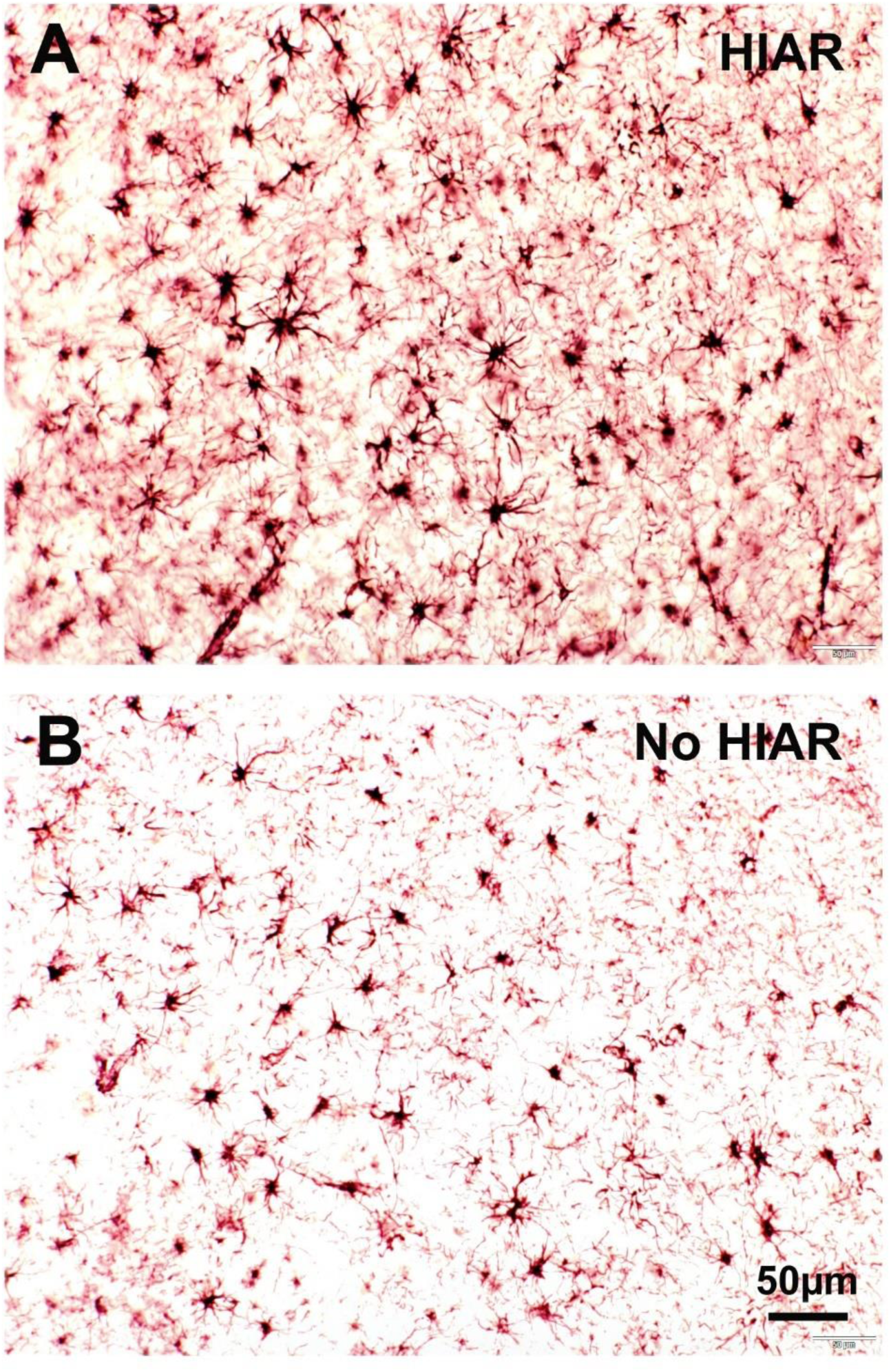
Expression of GFAP immunostaining in PFC of human brain post-fixed for 25 years. Following the treatment of citrate buffer (pH6.0) based heat-induced antigen retrieval (HIAR), abundant strongly stained GFAP-IR astrocytes were observed in both gray and white matter of PFC (**A**). However, without HIAR treatment, GFAP-IR astrocytes were less strong and less abundant (**B**) compared to that with HIAR treatment (**A**).

